# End-to-End Explainable AI: Derived Theory-of-Mind Fingerprints to Distinguish Between Autistic and Typically developing and Social Symptom Severity

**DOI:** 10.1101/2023.01.21.525016

**Authors:** Km Bhavna, Romi Banerjee, Dipanjan Roy

## Abstract

Theory-of-Mind (ToM) is an evolving ability that significantly impacts human learning and cognition. Early development of ToM ability allow one to comprehend other people’s aims and ambitions, as well as thinking that differs from one’s own. Autism Spectrum Disorder (ASD) is the prevalent pervasive neurodevelopmental disorder in which participants’ brains appeared to be marked by diffuse variations throughout large-scale brain systems made up of functionally connected but physically separated brain areas that got abnormalities in willed action, self-monitoring and monitoring the intents of others, often known as ToM. Although functional neuroimaging techniques have been widely used to establish the neural correlates implicated in ToM, the specific mechanisms still need to be clarified. The availability of current Big data and Artificial Intelligence (AI) frameworks paves the way for systematically identifying Autistics from typically developing by identifying neural correlates and connectome-based features to generate accurate classifications and predictions of socio-cognitive impairment. In this work, we develop an Ex-AI model that quantifies the common sources of variability in ToM brain regions between typically developing and ASD individuals. Our results identify a feature set on which the classification model can be trained to learn characteristics differences and classify ASD and TD ToM development more distinctly. This approach can also estimate heterogeneity within ASD ToM subtypes and their association with the symptom severity scores based on socio-cognitive impairments. Based on our proposed framework, we obtain an average accuracy of more than 90 % using Explainable ML (Ex-Ml) models and an average of 96 % classification accuracy using Explainable Deep Neural Network (Ex-DNN) models. Our findings identify three important sub-groups within ASD samples based on the key differences and heterogeneity in resting state ToM regions’ functional connectivity patterns and predictive of mild to severe atypical social cognition and communication deficits through early developmental stages.

## 1. Introduction

theory-of-Mind (TOM) is a major component of social cognition that allows an individual to attribute mental states to others and is also a key determinant of the quality of social interactions [1], [2]. Previous studies using functional MRI have found that the frontal-parietal network included the medial prefrontal cortex, posterior cingulate cortex, and bilateral temporoparietal junction (LTPJ and RTPJ) activated during the Theory-of-Mind tasks in typically developing [3], [4], [5]. One of the core deficits associated with Autism Spectrum Disorder (ASD) leads to genuine cognitive impairment in social cognition and communication deficits and mentalizing of concepts, which may particularly underlie deficits in Theory-of-Mind abilities [6], [7], [8], [9], [10], [11]. Despite decades of research, the neurobiology of ASD and core deficits in social cognition are still poorly understood. Few neuroimaging findings have been widely replicated, and no clear picture of the brain bases of socio-communicative and cognitive impairments in ASDs has emerged thus far [11]. Heterogeneity within the ASD Group is a significant challenge; individuals diagnosed with the same disorder can present with different behavioral symptoms. However, it is still tough to identify systematic neurological variation that correlates with the behavioral symptom that generalizes among neurotypical and Autism. There is also no clear roadmap to identify reliable and explainable neurobiological markers or behavioral symptom predictors, which can be helpful as accurate diagnostic measures. The availability of ‘Big Data’ and explainable artificial intelligence techniques provide contingency to identify ASD individuals and neurobiological features associated with ASD [12].

### 1.1 Related Work

In the past few years, DNNs have revolutionized the field of Ex-AI with major successes in applications such as computer vision, object recognition, natural scene processing, and natural language processing [12], [13]. DNNs, however, have had limited success in ab initio classification and identification of neurobiological features that distinguish neurodevlopmental and psychiatric disorders using functional brain imaging data [14]. This is due to several challenges in applying DNNs to brain imaging data, most notably dealing with the high dimensionality of whole-brain data and noisy measurements with a large degree of individual variability across data acquisition sites [14], [15]. A particular challenge here is the application to ASD, a neurodevelopmental disorder characterized by a spectrum of impairments and high levels of heterogeneity in phenotypic clinical symptoms [16], [17]. Past studies to classify ASD individuals from typically developing focusing on Theory-of-mind (ToM) brain regions using static functional connectivity (SFC) and BOLD time-series signals as feature sets as input to DNN and ML models did not always report self-consistent and generalizable findings [16], [17], [18], [19], [20], [21], [22], [23]. A few recent studies have attempted to use DNNs by reducing the dimensionality of brain data through explicit feature engineering [16], [18], [19]. Typically, in these approaches, precomputed static functional brain connectivities are provided as input to DNN models consisting of multiple fully connected networks followed by a sigmoid layer for classification. This approach has multiple problems. One of the key problems is that training DNN architectures with fully connected layers is challenging, particularly in neuroimaging applications, because of multiple free parameters and a limited number of labeled training data. As a result, these architectures tend to overfit the data and exhibit poor out-of-sample prediction [24]. Critically, extant approaches do not sufficiently exploit non-overlapping ToM and DMN region’s spatiotemporal characteristics with TD individuals and connectivity features, which contain more robust features of ASD phenotypes associated with self-related socio-cognitive impairments and mentalization. These non-overlapping ToM and DMN Spatio-temporal connectivity features after controlling for the variability are thought to inform on inter-individual anatomical, and phenotypic differences [**?**], [16], [25], [26].

### 1.2 Overview and Contributions of This Study

There are several novel aspects to this study. First, in the current study, we propose an end-to-end explainable and interpretable artificial intelligence-based computational model (refer Figure 1 and 2). The current study addresses the following key questions about neurodevelopemental differences in the heterogeneous profile of ToM abilities in children with ASD and the association between different levels of ToM development. To this end, we have developed an end-to-end explainable AI pipeline to identify a common source of variability that underlies the heterogeneous profile of ToM abilities between typically developing and ASD individuals focusing in particular during early development. Second, we show that the variability identified through the Contrastive Variational Autoencoder Model (CVAE) could be linked with non-clinical variables such as age, gender, and site-specific protocols. This allowed us to discover a distinct set of functional connectivity patterns in ToM regions in both groups and provided the best possible feature sets. Third, we show that ToM-DMN (understanding of others and self-related processing brain regions) connectivity features could be used as a set of biological features to train the classifiers to classify ASD and TD accurately. This finding is non-trivial as previous attempts lack explainable features of brain connectivity patterns in ToM regions and consistency and reliability to accurately predict their association with social symptom severity. We observed resting state hyper-connectivity within the brain regions anchored in ToM networks in the ASD group; in contrast, we found hypo-connectivity within the ToM regions in TD. Using the CVAE model, we have further shown that ASD-specific characteristics were correlated with clinical measures like ADOS-Total, ADOS-Social, FIQ, and functional connectivity. To the best of our knowledge, for the first time, proposed Ex-DNN and Ex-Ml models identified the ASD population from typically developing s by learning biologically interpretable connectivity features among ToM and DMN brain regions and achieved, on average, 95% accuracy with an F1 score of 0.94. The framework proposed here suggests that functional connectivity between the Left temporoparietal junction (LTPJ), Right temporoparietal junction (RTPJ), Right superior temporal sulcus(RSTS), Left angular gyrus(LAG), and precuneus(Prec) brain regions are the most robust feature set and may underlie the prevalence of heterogeneous profile of ToM abilities in children with ASD. These findings could be the first step in the association between brain connectivity fingerprints that link different ToM development levels.

**Fig. 1.**
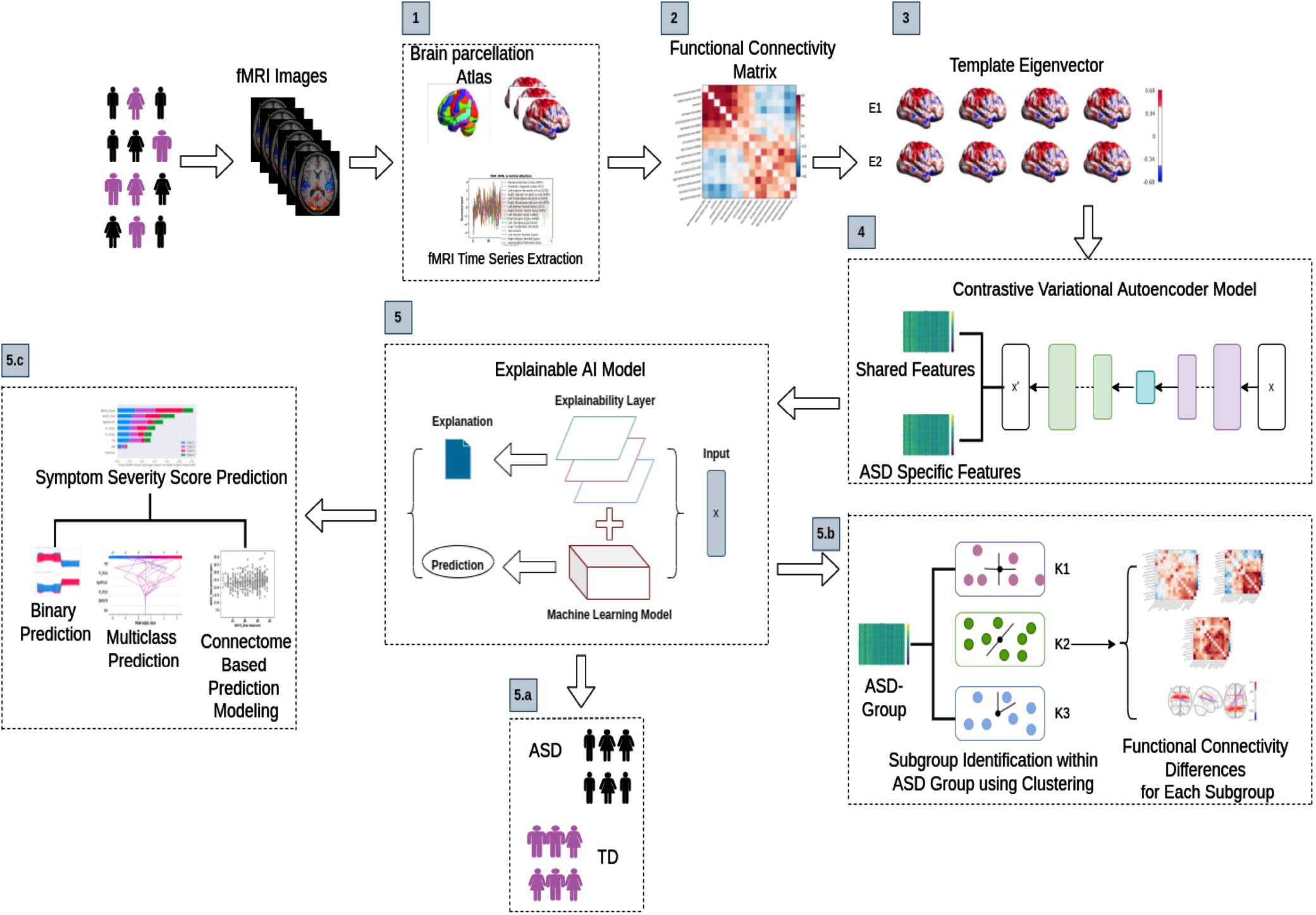
Illustrative overview of End-to-End Explainable Artificial Intelligence framework for identifying the common source of variability that helps classify ASD samples from typically developing using Theory-of-Mind brain region and identification of heterogeneity or sub-groups within ASD samples and their symptom severity score prediction. The contribution of the paper is as follows: step 1 is data extraction; steps 2 and 3 are the calculation of the Functional Connectivity matrix and its low-dimensional representation; step 4 is the identification of the source of variability; step 5 is a classification of ASD group: Steps 5. a and 5. b is the identification of the ASD subgroup and their symptom severity score prediction.

**Fig. 2.**
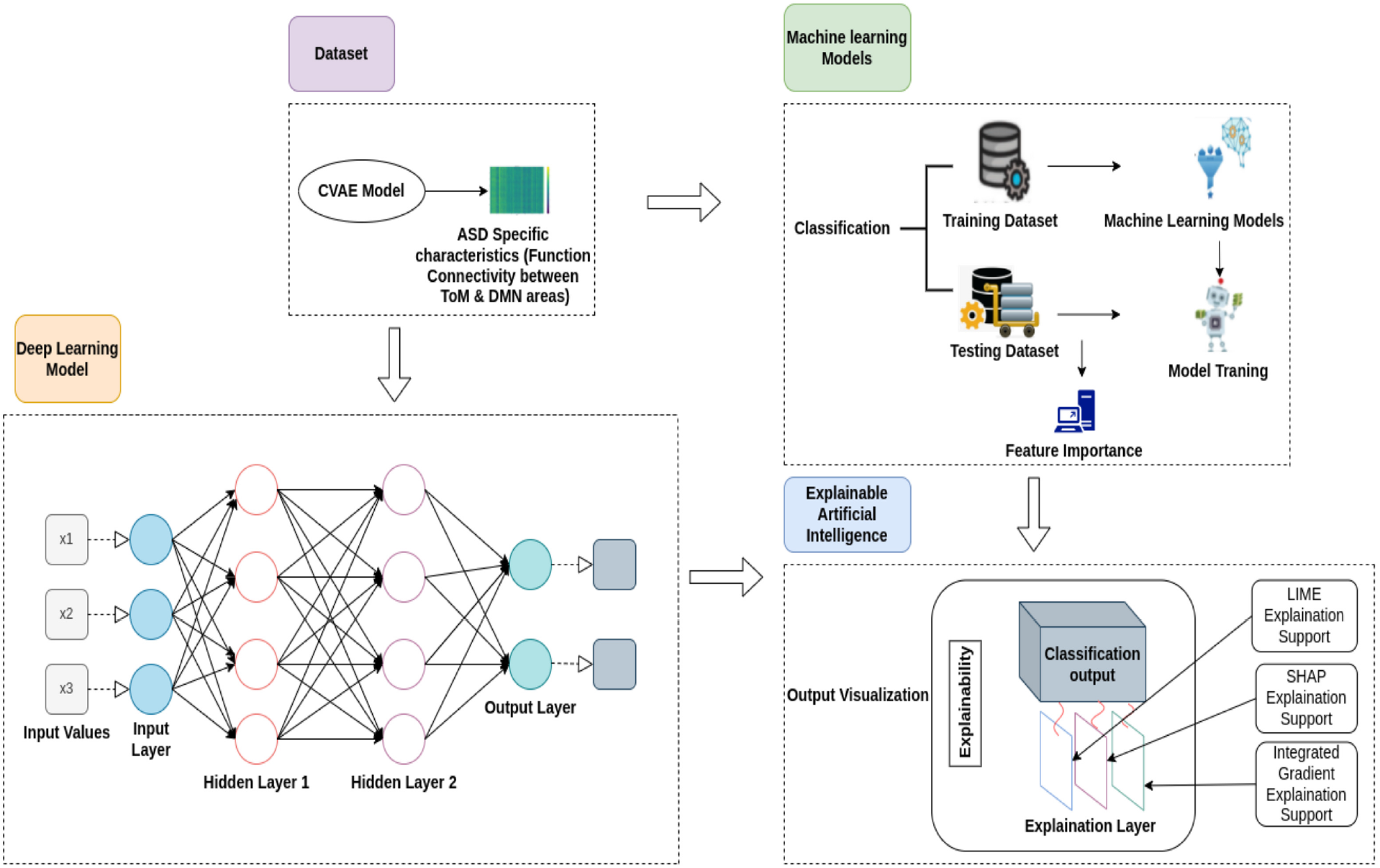
Architecture of Explainable AI model. The characteristics (resting state Functional Connectivity) identified by the CVAE model are provided to DNN and ML models for classifying ASD from typically developing. The integrated Gradient, LIME, and SHAP layers are applied to determine black box brain features that are associated with ASD and underlie classification

Our study systematically addresses and overcomes several key challenges and previous limitations: a) The first limitation that we address here is the site-specific heterogeneity while using consortium data. The data cannot be expected to comprise from a single site as that offers only a limited sample, posing a challenge for developing and training EAI models. Hence, there is a requirement to acquire and pool images across multiple sites, which can introduce significant heterogeneity due to engineering sampling bias and biological sampling bias. We overcome this challenge by proposing a Combat harmonization technique based on empirical Bayes methods. [27]. Our method handles heterogeneous data from different imaging protocols while learning robust representations to classify and identify robust neurobiologically meaningful features that distinguish individuals with ASD. b) The second major issue is selecting Brain regions associated with Theory-of-Mind (ToM). We have selected the Regions-of-Interest (ROI) from the Human Connectome Project (HCP) available on NeuroVault and extracted Bold time-series signals from TOM and Default Mode Network (DMN) areas [5]. The reason behind using Default-mode Network alongside the ToM network is based on the existing literature suggesting default-mode network (DMN) nodes would feature prominently in both classification and symptom prediction. Many of these areas show specific differences between autistic and TD, in that DMN node would feature prominently in ToM and mentalization concept in both classification and symptom prediction [5].

c) The third limitation is identifying a common source of variability in typically developing s and the autistic using Theory-of-Mind brain regions, which may confound our findings based on the neuroimaging and connectivity-derived phenotypes. To identify ASD-Specific variation, we segregate it from common variation between typically developing and Autistic using a Contrastive-variational Autoencoder model that can distinguish clinically applicable individual variation. It segregates its internal architecture into ASD-specific and shared features.

d) The fourth limitation is the classification of ASD from typically developing s. The proceeding step was able to identify ASD-specific and shared characteristics. We have implemented an Explainable AI model that can accurately classify ASD samples from typically developing s and identify neurobiological or brain features responsible for classification without biasing engineering features. We have used Integrated Gradient, LIME(Local Interpretable Model Agnostic Explanation), and SHAP(SHapley Additive exPlanations) approaches to identify features that underlie the classification.

e) The fifth limitation is to identify heterogeneity within the ASD group. To capture variability to identify the sub-groups of ASD samples, we have implemented a k-means clustering algorithm, and the no. of clusters is determined using the elbow approach. Using this approach, we are getting 3 clusters. We have also calculated functional connectivity differences between all the sub-groups to check whether there is any overlapping in connectivity.

f) The sixth limitation is identifying symptom severity score predictions associated with the heterogeneous demonstration of ASD. The major issue we address here is identifying neurobiologically interpretable and meaningful features rather than requiring feature engineering for predicting ASD social and communication deficits with heterogeneous manifestations in ASD children arising from atypicality of TOM and DMN regions functional connectivity patterns. To the best of our knowledge, no previous deep-learning classification study has investigated brain features that robustly predict TOM clinical symptoms without feature engineering.

We illustrate that our ExAI approach (Refer Figure 1): a) accurately identifies the source of variability in ASD and TD, b) Classify ASD samples from TD, and also identifies the brain features that underlie the classification, c) Identifies heterogeneity that exists within ASD samples, d) This approach also determines ToM brain features that can predict symptom severity scores of each ASD sub-group.

Section 2 describes the participants’ details, fMRI acquisition protocols, and Preprocessing methods employed, Next, we describe the selection of ToM brain regions and estimation of static functional connectivity (SFC) in ToM brain regions using ASD and TD resting state BOLD fMRI signals and a low-dimensional representation of SFC. In section 3, we disentangle common variations between TD and ASD participants using a Contrastive-Variational Autoen-coder model(CVAE) that allowed us to distinguish clinically associated individual variations; subsequently, In section 4, we implement multiple Explainable DNN/ML models where we provided resting-state FC matrix from ToM and DMN brain areas as feature set for classifying ASD and TD individuals, Finally, we identify a set of neurobiological features underlying classification performance in our proposed model which allow us to discover subgroups within ASD participants and relate their ToM-DMN functional connectivity and prediction of Autistic symptom severity.

## 2 Study cohort for characterizing ASD-specific variations

### 2.1 Participants and MRI Preprocessing

In the current study, we analyzed ABIDE I dataset that underwent preprocessing and quality control. Finally, it comprised 400 Autism samples and 460 TD samples (Total No. of samples = 860). All the neuroimaging data were preprocessed using Configurable Pipeline for the Analysis of Connectomes (C-PAC), which included slice-time correction, motion correction, functional normalization, and smoothing. ABIDE dataset included fMRI data acquired from multiple sites, which resulted in the problem of heterogeneity due to Engineering sampling bias and Biological sampling bias. We addressed this challenge using the Combat harmonization technique: Assuming that the errors induced in the imaging characteristics may be standardized by altering the location (means) and scale (variances) across the batches. We performed harmonization using multivariate linear mixed-effects regression. This method used empirical Bayes for the improvement of the estimation of the model. [27], [28], [29], [30]. It removed unwanted variation associated with the site and maintained biological association in the dataset.

### 2.2 Resting-State Functional Connectivity

As individuals with ASD display extensive and robust deficits in ToM, we extracted BOLD time series signals from Theory-of-Mind and other large-scale brain networks associated with social cognition, i.e., Default Mode Network(DMN). The Regions-of-Interest(ROI) for this purpose was derived from Human Connectome Project(HCP) available on Neuro Vault [5]. We used the Harvard-oxford atlas and created ROIs (that included the Medial prefrontal Cortex (MPFC), Posterior Cingulate cortex (PCC), Left superior temporal sulcus (LSTS), Right superior temporal sulcus (RSTS), Left Temporoparietal Junction (LTPJ), Right Temporoparietal Junction (RTPJ), Left Inferior Frontal Gyrus (LIFG), Right Inferior Frontal Gyrus (RIFG), Left Angular Gyrus (LANG), Right Angular Gyrus (RANG), Left Cerebellum (LCEREB), Right Cerebellum (RCEREB), Precuneous (PREC), Left Inferior Parietal Cortex(LIPC), Right Inferior Parietal Cortex(RIPC), and Ventrolateral Prefrontal cortex(VLPFC))using a spherical binary mask with a 10 mm radius (see Table 2 of supplementary material). The average time series BOLD signals for each participant were extracted for Pearson’s correlation analysis using the following formula [31]:

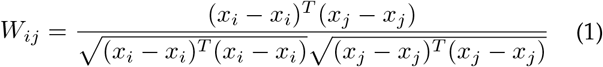

Where *x*_*i*_ ∈ *R*^*t*^ is the time series associated with defined brain regions. t is time node, i = 1,2,…….,n, where n is number of ROIs. After that, using an inverse hyperbolic inverse function, correlation coefficients were Fisher’s z-transformed. We also applied false discovery rate (FDR) correction for each connectivity analysis (see Supplementary material section 1 for details).

**Table 1:** Demographic Information of ASD and TD groups From ABIDE dataset.

| Sr. | Age (In Years) | Gender( Male/Female ) | IQ | ADI-R Social | ADI-R Verbal |
| --- | --- | --- | --- | --- | --- |
| ASD- group (400 Samples) | 13.9 ± 6.6 (range: 6 to 42) | 360\40 | 105 ± 17 | 18.8 ± 5.5 | 15.4 ± 4 .5 |
| TD-Group (460 Samples) | 14.9±5.9 (range: 6 to 40) | 370\90 | 112 ± 14 | | |

**Table 2:** Brain region with MNI coordinates that are associated with Theory-of-Mind concept.

| Sr. No. | Region-of-Interest | MNI-Coordinates(X,Y,Z) |
| --- | --- | --- |
| 1 | Medial prefrontal Cortex (MPFC) | 0, 48, -18 |
| 2 | Posterior Cingulate cortex (PCC) | 0, -52, 22 |
| 3 | Left superior temporal sulcus (LSTS) | -58, -48, 7 |
| 4 | Right superior temporal sulcus (RSTS) | 58, -44, 8 |
| 5 | Left Temporoparietal Junction (LTPJ) | -58, -58, 22 |
| 6 | Right Temporoparietal Junction (RTPJ) | 58, -52, 20 |
| 7 | Left Inferior Frontal Gyrus (LIFG) | -48, 14, 26 |
| 8 | Right Inferior Frontal Gyrus (RIFG) | 48, 14, 26 |
| 9 | Left Angular Gyrus (LANG) | -46, -72, 31 |
| 10 | Right Angular Gyrus (RANG) | 48, -69, 31 |
| 11 | Left Cerebellum (LCEREB) | -20, -72, -36 |
| 12 | Right Cerebellum (RCEREB) | 20, -72, -36 |
| 13 | Precuneous | 0, -49, 40 |
| 14 | Left Inferior Parietal Cortex | -45, -46, 53 |
| 15 | Right Inferior Parietal Cortex | 52, -42, 50 |
| 16 | Ventrolateral Prefrontal cortex | 42, 46, 0 |

### 2.3 Low-Dimensional Representation of Functional Connectivity

We calculated the static functional connectivity (FC) matrix from resting-state fMRI data. We finally generated a low-dimensional representation of the FC matrix based on the Eigen Decomposition method. We implemented the Principal component dimensionality reduction technique in which each eigenvector was aligned with the template manifold and measured using group-averaged functional connectivity. We selected two principal eigenvector components that capture the variability of data without losing essential biological features and explained approximately 68 % of the information of the template FC matrix. Using PCA, we identify eigenvectors that can be provided as input to the next step to check whether resting-state SFC is associated with ASD or TD characteristics.

### 2.4 Identification of Shared and Specific Variability

To characterize ASD-Specific variation, we disentangle it from common variation between typically developing and Autistic individuals using a Contrastive-Variational Autoen-coder model(CVAE) that allowed us to distinguish clinically applicable individual variation [26]. It dissociated its internal architecture into ASD-specific and shared characteristics. CVAE contained of an encoder and decoder, in which the encoder included 2 consecutive convolutional layers with stride: 2, kernel size: 3, and convolutional filters: 64 and 128 for shared characteristics and 2 convolutional layers with the same parameters for ASD-specific characteristics [26]. Instead of projecting data into one latent space, this encoder projected data into two separate 16-dimensional latent distributions: 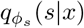 and 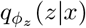 [32], where s and z were latent variables. We used the following lower bound likelihood for shared characteristics [32]:

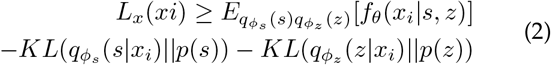

For lower bound likelihood for ASD-specific characteristics:

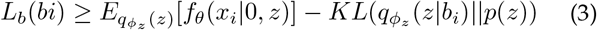

where 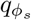 and 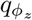 are two encoder network and *f*_*θ*_(.) is single decoder network. The decoder used 2 de-convolutional layers with convolutional filters: 64 and 128, that concatenated the two latent distributions into a single 32-dimensional vector and accepted it as input to produce output.

We applied Principal Component Analysis (PCA) to the FC matrix to get a low-dimensional matrix representation and extracted 2 eigenvectors. We provided eigenvectors belonging to other clinical and non-clinical parameters to the CVAE model as input to check whether it is associated with a shared or ASD-specific variation. We used Representational Similarity Analysis(RSA) approach to compare CVAE’s ASD-specific and shared characteristics for each individual (see Supplementary material section 1 for details). As a result, shared characteristics were associated with non-clinical measures, whereas ASD-specific characteristics were associated with clinical measures and resting-state Functional Connectivity.

## 3 Cross-validation and classification Analysis of ABIDE Cohort Data

### 3.1 Classification Using Explainable AI and Analysis Using Five-Fold Cross Validation

After getting ASD-specific characteristics, we implemented multiple Explainable DNN/ML models where we provided resting-state FC matrix from ToM and DMN brain areas as a feature set for classifying ASD and typically developing individuals. The main idea behind using Explainable AI models was to know which brain region’s connectivity acted as predictors in machine learning models and their comparative importance (Refer to Figure 2). We developed an interpretable Explainable AI architecture, in which we applied multiple machine learning and deep learning models with explanatory support of LIME(Local Interpretable Model Agnostic Explanation), SHAP(SHapley Additive exPlanations), and Integrated Gradient (IG). For classification using Deep Neural network (DNN), we used the Adam algorithm to train and test the DNN model with three layers in which we provided ASD-specific features to first layer and shared features to the second layer that contained 300 neurons in each layer (i.e., Patience = 3, Metric = validation loss) and applied the Relu activation function and finally the final output layer comprised of a conventional softmax function for classification. The purpose behind using the Adam algorithm was that it was able to combine the best features of the Adaptive Gradient Descent (AdaGrad) and Root Mean Square (RMS) Prop algorithm to provide an optimization algorithm. Instead of accommodating learning rates based on the mean (i.e., first moment), it also considered the gradient’s uncentred variance (second moment). We implemented Adam algorithm using the following formula;

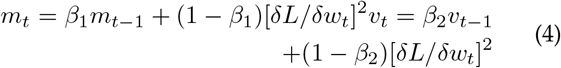

Where *β*_1_ and *β*_2_ are decay rates of an average gradient. For classification using the machine learning algorithm, we implemented multiple ML algorithms. We trained and tested the models on the same ASD-specific and shared features. The essential advantage of our approach was that it did not require any spatiotemporal dynamics features; it could give the best performance in classifying ASD and typically developing only using SFC between Theory-of-Mind, and Default-mode-network brain areas and providing parsimonious brain features underlying classification. To reduce the chance of bias and report low variance, we implemented five-fold cross-validation and a Leave-one-out method to evaluate the model’s performance (precision, recall, accuracy, F1-score) (see Supplementary material section 1 for details).

### 3.2 Identification of neurobiological features underlying classification

We used LIME(Local Interpretable Model Agnostic Explanation) and SHAP(SHapley Additive exPlanations), and an Integrated Gradient-based feature diagnostic approach to identify neurological features that were responsible for classification.

To identify the features responsible for the classification, we implemented an Integrated Gradient(IG) for the interpretability of the DNN model, which computed the gradient for the output predicted by the model to its input feature. IG selected a baseline to produce a high entropy prediction that described uncertainty, then calculated feature attribution associated with an ambiguous baseline. Finally, IG interpolated the baseline with the prediction from uncertainty to certainty towards actual input. We implemented IG using the following formula:

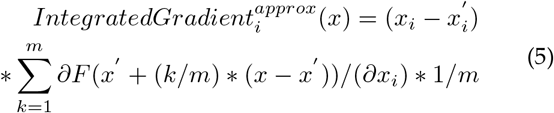

Where: Input features = (*∂F* (*x*′ + *k/m* * (*x* − *x*′)))*/*(*∂x*_*i*_), Average gradients 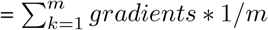

Because of the limitations of IG toward machine learning, we implemented a) LIME(Local Interpretable Model Agnostic Explanation) based on learning an explainable or interpretable model locally throughout the prediction using the following formula:

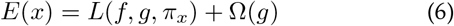

Where: Ω(*g*) is the complexity of the explanation of the model, *π*_*x*_ is locality. b) SHAP(SHapley Additive exPlanations) that decomposed the final output prediction to identify the contribution of each attribute for the explainability of machine learning models using the following equation:

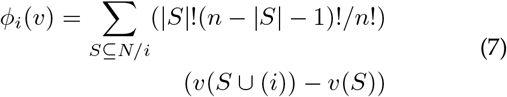

Where: n is total no. of regions, N contains all possible subsets, |*S*| no. of features in S, and v(s) calculated contribution of S.

### 3.3 Identification of Sub-Group Within ASD Group

The ASD sample’s resting-state FC was subjected to an unsupervised K-means algorithm to identify heterogeneity or subgroups within ASD samples. We implemented the K-means clustering algorithm and got three clusters using the elbow approach (see Supplementary material section 1 for details) [33]. We also implemented a variational auto-encoder model to validate the results and provided resting-state FC as input. We used latent space as a feature set and applied the K-means clustering algorithm to find sub-groups.

### 3.4 Functional Connectivity Differences in each Sub-Group

To identify Functional Connectivity differences, We extracted average time series BOLD signals from Theory-of-Mind and Default-Mode Network brain areas for each individual in every cluster and performed Pearson’s correlation analysis. We calculated the Functional connectivity matrix for each cluster and performed multiple t-tests with a p-value*<*0.01. We have finally performed false discovery Rate (FDR) correction for every connectivity. We also conducted a matrix similarity analysis to check if connectivity was overlapping among the three clusters. For that, firstly, we extracted the top or upper triangle of the functional connectivity matrix as it provided more consistency with the matrix function and performed spearman correlation for each combination of the upper matrix. To validate the results, we performed permutation for test significance. For each iteration, we stumbled over the sequence of rows and columns from one of the correlation matrices and again calculated the similarity between the two matrices. Here, the amount of time for permutation was equal to 5000 times. Finally, we checked how many values survived after this test to obtain a p-value*<*0.05.

### 3.5 Symptom Severity Score Predictions in ASD Using Multiple Approaches

We investigated the association between neurobiological features identified by Explainable AI models and symptom severity scores (see Supplementary material section 1 for details). We investigated whether resting state functional connectivity could identify ASD-symptom severity scores for each individual. We implemented multiple approaches for symptom severity scores prediction like Binary prediction, Multiclass prediction, and Connectome-Based modelling Prediction(CPM) [34]. We implemented Binary prediction using machine learning models like regression, which was a very basic approach and very limited (it applied to identify whether the participant is severe ASD or not). Because of the limitation of binary prediction, we moved towards multi-class prediction, in which we created four classes of severity (low, moderate, high, and very high). We performed this using a Deep learning model with a sequential optimizer with three layers (loss function = Binary cross-entropy) and applied the sigmoid activation function; we also used SHAP to interpret this model. We also implemented multiple machine learning algorithms and an ensemble learning approach with explainability for multi-class prediction. To predict each individual’s ADOS-Total and ADOS-social scores, we implemented Connectome-based prediction modeling, which used a functional connectivity matrix extracted earlier and fitted a linear model for the brain-behavior relationship. This model was able to predict scores for each individual.

## 4 RESULTS

### 4.1 Extraction of Time-Series Signals Specific to Theory-of-Mind Areas and Low-Dimensional Representation of Functional Connectivity

We extracted time-series signals from Theory-of-Mind, and Default-mode Network brain areas and constructed a 16*16 static functional connectivity matrix by calculating Pearson’s correlation. To validate the results, we performed multiple one-sample t-tests with a p-value *<* 0.01 and applied FDR correction. For low-dimension representation, we used the principal component analysis method for extracting the top two principal components that contained approximately 68 % of the information.

### 4.2 Identification of Shared And ASD-Specific Features or Source of Variability

To identify the variability between ASD and TD, we first implemented the Variational Autoencoder model(VAE), which allowed us to check whether VAE could identify an association between ASD and TD without disassociating ASD-specific and shared variation; for that purpose, we used Representational Similarity Analysis(RSA), in which we first calculated a pair-wise dissimilarity between participants for VAE typically developing features and acquired a dissimilarity matrix. We repeated this process for each clinical and non-clinical characteristic. Finally, using the Kendall correlation coefficient, we compared the VAE dissimilarity matrix with the matrix for every individual (Refer Figure 3A). As a result, result from VAE showed Kendall correlation with some non-clinical characteristics (ScannerID (*τ* = 0.01, t(9)=0.74, p-value *<* 0.042), Gender (*τ* = 0.01, t(9)=1.94, p-value *<* 0.04)) and also correlated with DSM-IV(Diagnostic Statistical Manual IV)(*τ* = 0.02, t(9)=0.54, p-value *<* 0.03) but did not get any correlation with ADOS Total, ADOS Social, FIQ, and functional Connectivity. Hence, the VAE model failed to encapsulate variation in characteristics (see Supplementary material section 2 for details).

**Fig. 3.**
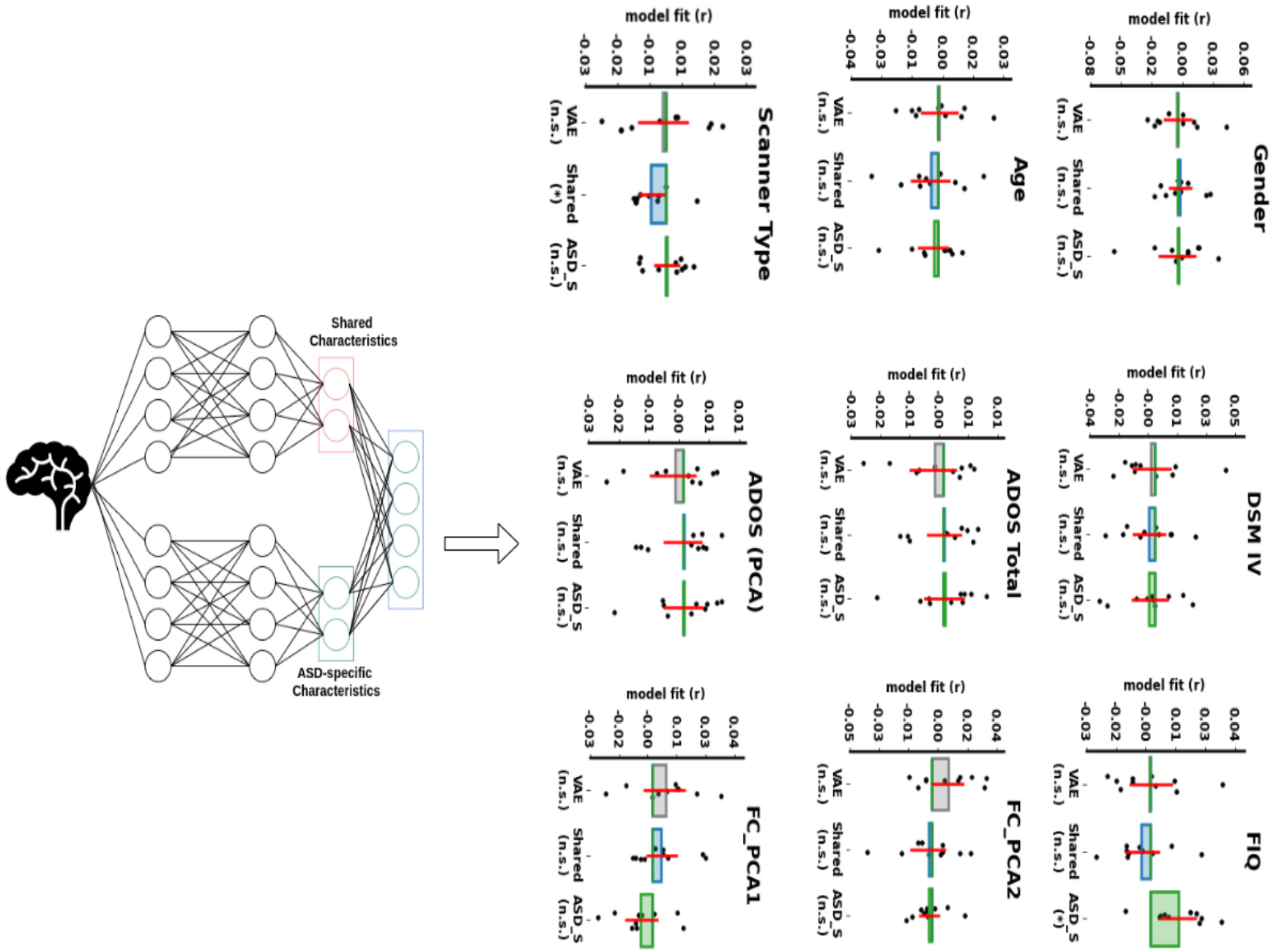
Comparison of Variational and Contrastive-variational Autoencoder model’s matrices. CVAE features best capture the common variation between ASD and TD groups, and parameters associated with ASD are best obtained by clinically ASD-specific measures and resting-state functional connectivity. VAE model is performing worst in all conditions.

To overcome the limitation of VAE, we implemented the Contrastive Variational Autoencoder model(CVAE), which segregated its interior characterization into ASD-specific and shared variations, which allowed us to identify clinically related individual variations. To compare CVAE’s ASD-specific and neuro-typical characteristics with every individual variation, we again used RSA (Refer Figure 3A). As a result, we found that shared characteristics were correlated with non-clinical characteristics (i.e., ScannerId (*τ* = 0.01, t(9)=0.85, p-value *<* 0.0.39), Age (*τ* = 0.03, t(9)=1.41, p-value *<* 0.04), Gender (*τ* = 0.02, t(9)=0.51, p-value *<* 0.045)); in contrast, ASD-specific characteristics were associated with clinical characteristics (i.e., ADOS Total (*τ* = 0.04, t(9)=2.65, p-value *<* 0.03), ADOS Social (*τ* = 0.04, t(9)=1.88, p-value *<* 0.01), FIQ (*τ* = 0.02, t(9)=1.95, p-value *<* 0.04)), as well as resting-state Functional Connectivity (*τ* = 0.04, t(9)=1.32, p-value *<* 0.02).

### 4.3 Classification of ASD and TD Individuals Using Explainable AI

To classify ASD individuals from typically developing individuals, we implemented multiple explainable machine learning models and explainable deep learning models. The results from the CVAE model suggested that resting-state FC was associated with ASD-specific characteristics. First, we trained our explainable ML/DNN models using resting-state functional connectivity on ABIDE dataset with a ratio of 70:30. ExML models achieved an average accuracy of 90 % with an F1 score of 0.91, with a sensitivity of 0.87 and specificity of 0.85. In contrast, the ExDNN model achieved an average accuracy of 96 % with an F1 score of 0.95, sensitivity of 0.95, and specificity of 0.93. We implemented Five-fold cross-validation and the Leave-one-out approach to validate the model’s performance and achieved an average accuracy of 96 % across all five-fold cross-validation and leave-one-out methods (see Tables 3 and 4 from supplementary material). The essential advantage of this approach was that it used only static functional connectivity from the brain regions anchored in only two major brain networks implicated in ToM processing namely Theory-of-Mind, and Default-mode Networks (see Supplementary material section 2 for details).

**Table 3:**
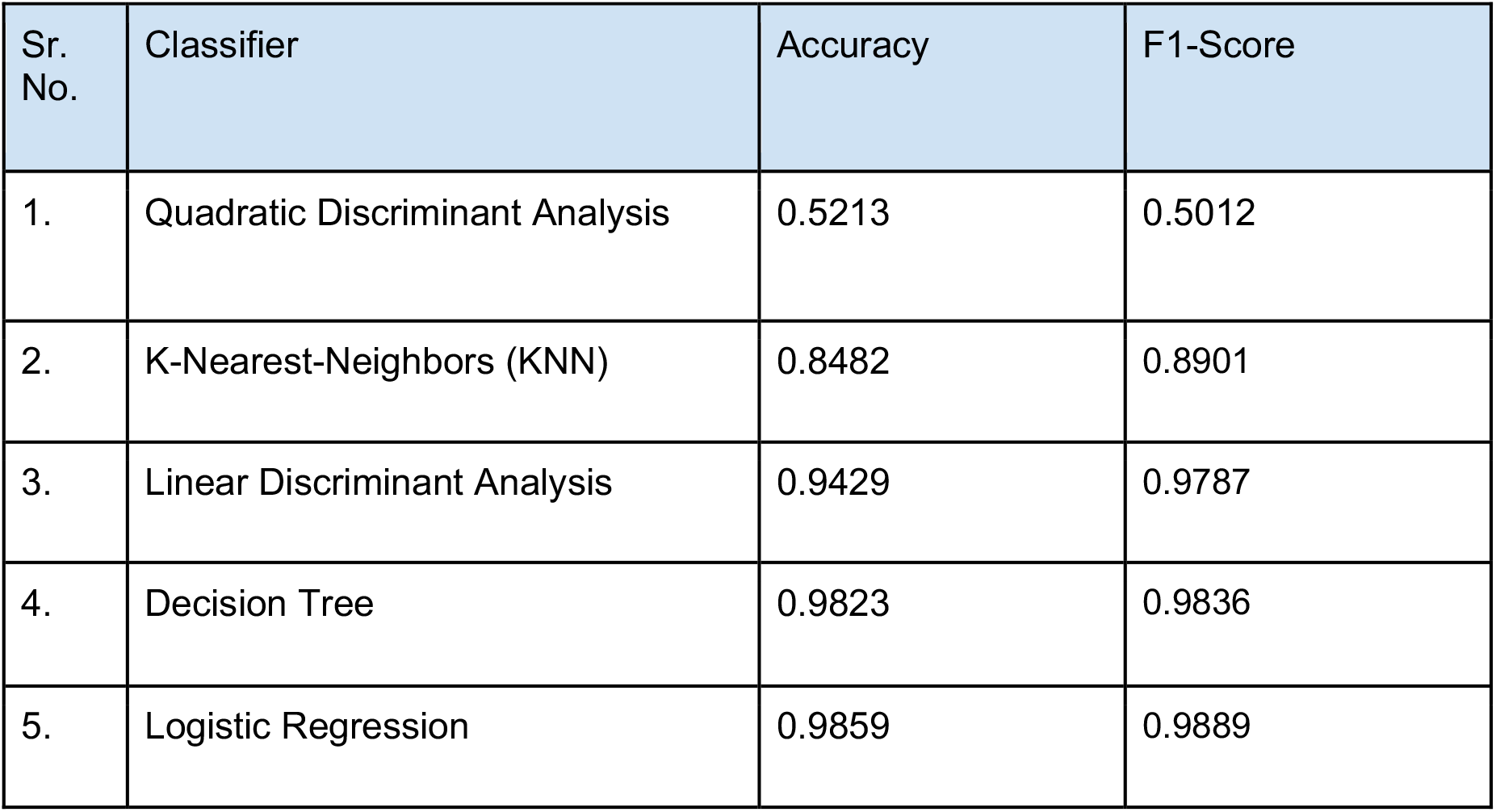
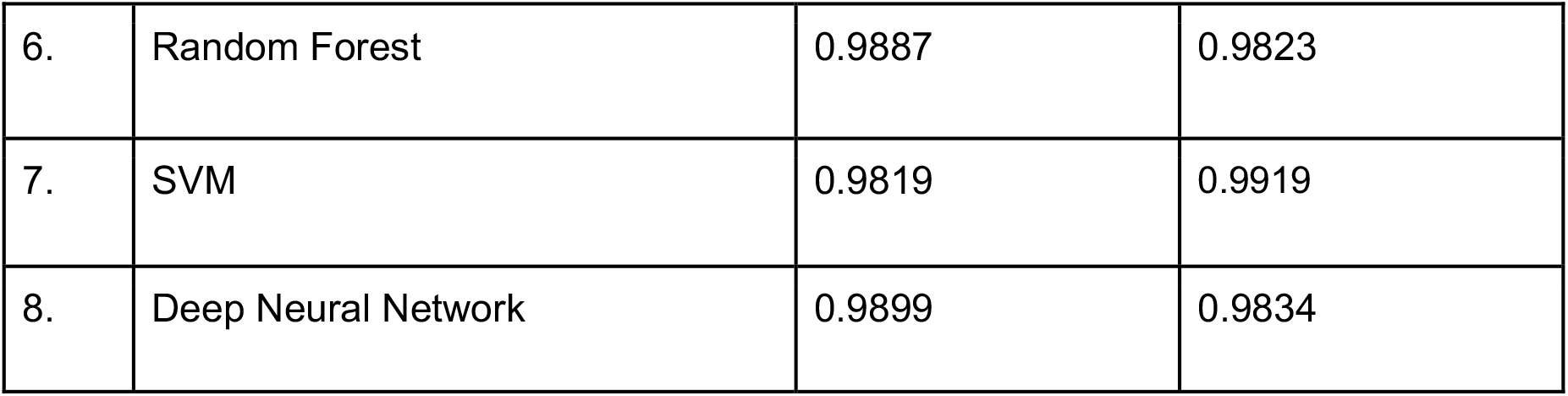
ASD vs. TD classification accuracies on ABIDE Cohort using ASD-Specific feature set.

**Table 4:** ASD vs. TD classification accuracies using five-fold cross-validation on ABIDE cohort.

| Sr. No. | Classifier | Accuracy | F1-Score |
| --- | --- | --- | --- |
| 1. | Quadratic Discriminant Analysis | 0.5912 | 0.5012 |
| 2. | K-Nearest-Neighbors (KNN) | 0.8567 | 0.8901 |
| 3. | Linear Discriminant Analysis | 0.9624 | 0.9787 |
| 4. | Decision Tree | 0.9831 | 0.9836 |
| 5. | Logistic Regression | 0.9913 | 0.9889 |
| 6. | Random Forest | 0.9865 | 0.9823 |
| 7. | SVM | 0.9876 | 0.9919 |
| 8. | Deep Neural Network | 0.9901 | 0.9934 |

For explainability, we applied LIME(Local Interpretable Model Agnostic Explanation), SHAP(Shapley Additive ex-Planations), and Integrated Gradient-based approaches, which provided the information on which extent each input feature contributed to the classification (Refer Figure 3B, and 3C). We computed the median of feature scores and identified ROIs that contributed 5% in classification. We observed that connectivity between the Left temporoparietal junction, Right temporoparietal junction, Right superior temporal sulcus, Left angular gyrus, Posterior cingulate cortex, and precuneus contributed most to classification. These biological features can distinctly demarcate ASD from typically developing individuals.

### 4.4 Identification of Sub-Groups Within ASD samples and Functional Connectivity Differences in Each Sub-Group

After getting the ASD samples, we applied K-means clustering to find sub-groups within ASD samples. Using the elbow approach, we obtained three clusters of different sizes (Cluster 1 contained 185 participants, cluster 2 had 98, and cluster 3 contained 117) (see supplementary material section 1 for details). For validating results, we implemented the variational autoencoder model and K-means on results coming out from the variational autoencoder model and again got 3 clusters. To identify functional connectivity differences between all clusters, we extracted time-series BOLD signals from Theory-of-Mind and Default-mode Network brain areas (Refer Figure 4). Finally, we calculated the Pearson correlation coefficient to obtain a functional connectivity matrix. We implemented multiple one-sample t-tests with p-value *<*0.01 and FDR correction on each connectivity for validation. Cluster 1 was close to the typically developing group, i.e., with the lowest ASD symptom severity scores, whereas clusters 2 and 3 were strongly associated with ASD-symptom.

**Fig. 4.**
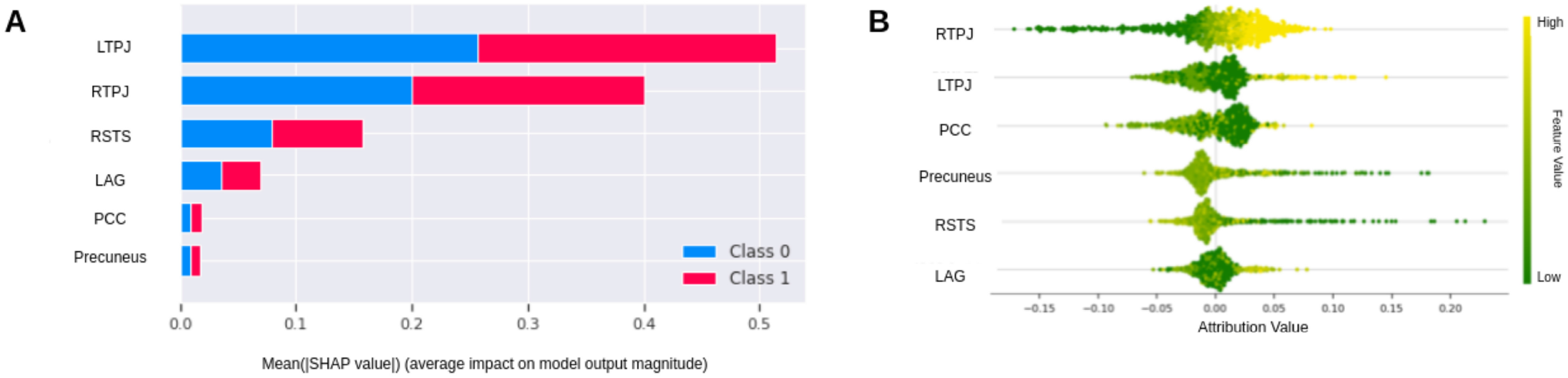
(A) Importance of Features that underlie classification for ASD and TD group in Explainable Machine learning model using LIME and SHAP. (B) Importance of Features that underlie classification for ASD and TD group using Explainable Deep Neural Networks using Integrated gradient.

**Fig. 5.**
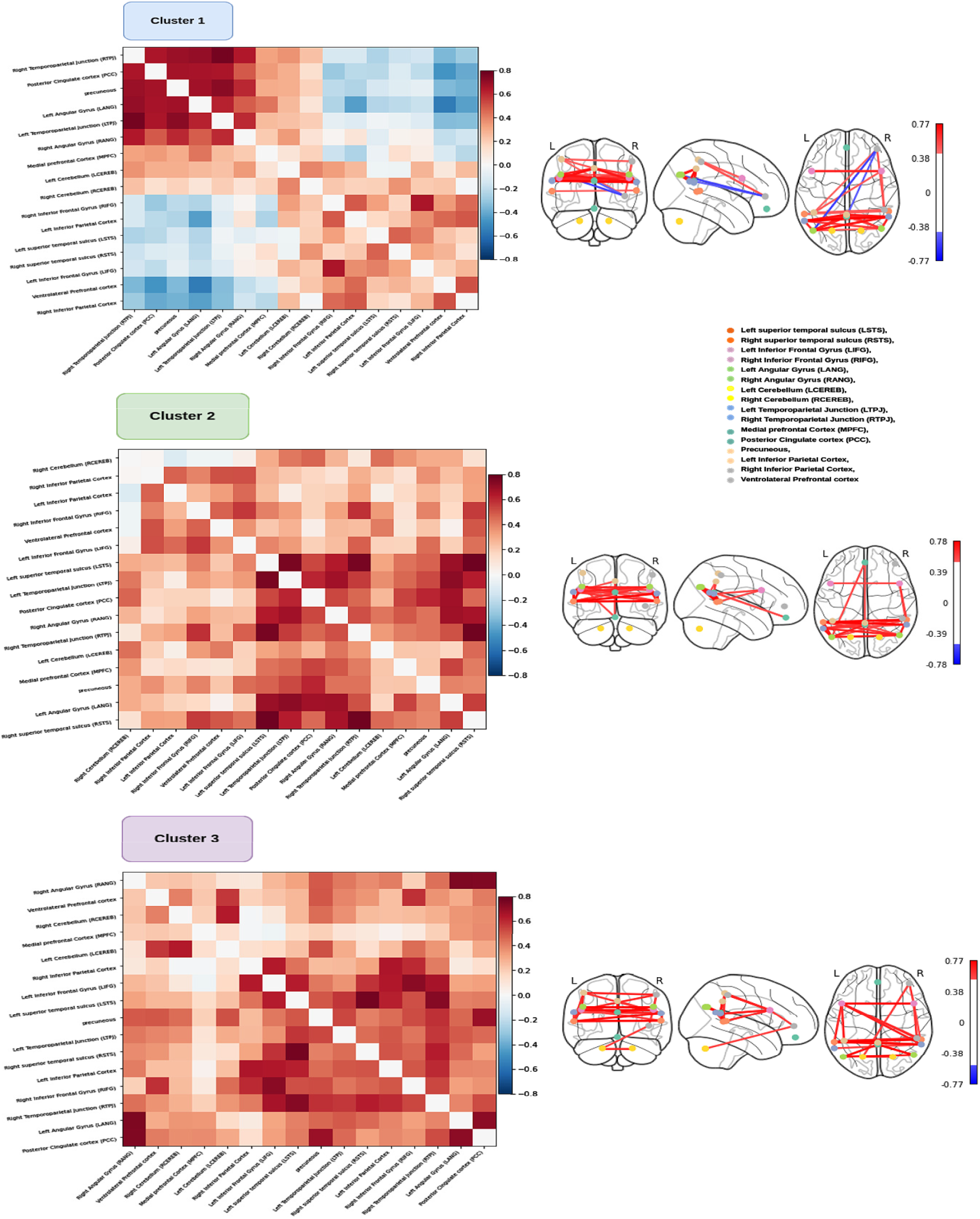
Cluster 1: First sub-group and its functional connectivity matrix within Theory-of-Mind and default-mode Network brain areas that is present within ASD Group. It showed that this cluster is not that severe to ASD. Cluster 2: Second sub-group and its functional connectivity matrix within Theory-of-Mind and Default-mode Network brain areas. This sub-group is associated with ASD severity. Cluster 3: Third sub-group and its functional connectivity matrix within theory-of-Mind and Default-mode Network brain areas. This sub-group is also associated with ASD severity.

We also conducted a matrix similarity analysis to check if connectivity is overlapping among the three clusters (Refer Figure 4). First, we extracted the upper triangle of the functional connectivity matrix and performed a spearman correlation for each combination of the upper triangular matrix. To validate the results, we performed permutation in which, for each iteration, we shuffled the sequence of rows and columns from one of the correlation matrices and again calculated the similarity between the two matrices. Here, we performed permutation 5000 times. Finally, we checked how many values survived after this test to obtain a p-value *<*0.05.

### 4.5 Symptom-Severity Score Prediction

After the steps described in sections 4.1-4.4, we investigated whether resting-state functional connectivity of ToM and DMN areas could identify ASD-symptom severity scores for each individual. We applied multiple approaches, i.e., Binary prediction, Multiclass prediction, and Connectome-Based prediction modelling. In Binary prediction, we applied multiple Explainable DNN/ML models to predict the severity of individuals (see supplementary material section 2 for details). In Multiclass prediction, we divided ASD-specific severity into four severity classes: Low, moderate, high, and very high. Then we applied multiple Explainable Machine learning and deep learning algorithms (Refer Figure 6). We got the best accuracy of 78 % with an F1-score of 0.78 from the Explainable Ensemble learning approach and 75 % with an F1-score of 0.73 accuracies using DNN (see Tables 5 and 6 in the supplementary material). After that, we applied CPM based on linear-regression models for extracting and encapsulating the most appropriate features from fMRI time series signals to predict the behavioral scores of each individual. We found that the CPM approach could predict ADOS-Social scores but not ADOS total scores using ToM and DMN brain areas functional connectivity. We found CPM a convenient and suitable technique for behavioral score prediction and fingerprinting, which could predict ADOS-Social scores accurately for each individual (refer to Supplementary material section 2 for more details).

**Fig. 6.**
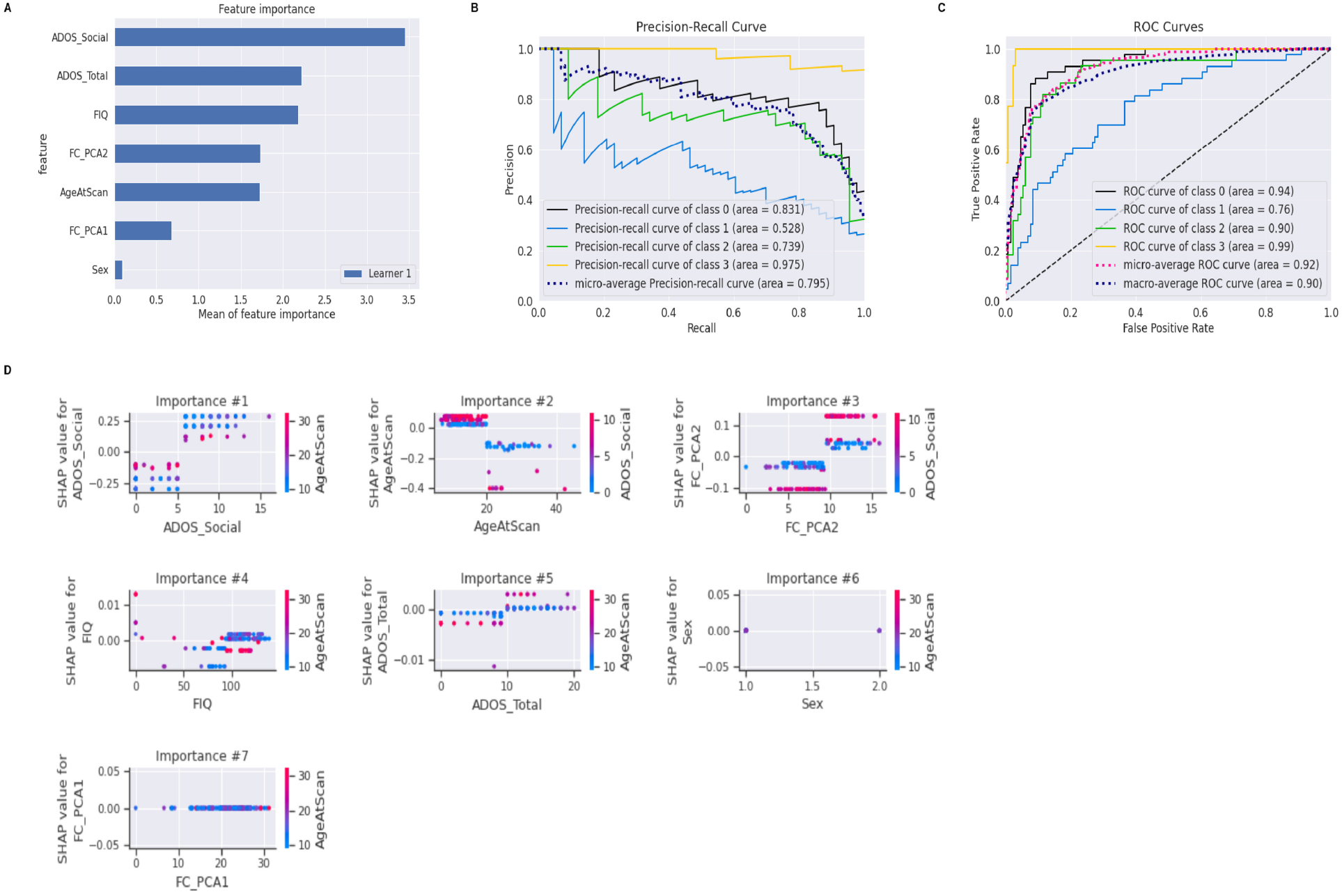
Prediction of severity to ASD using Multiclass Prediction. A) Identification of features that are responsible for the prediction of classes of severity which is showing that ADOS-Total and ADOS-Social contributed the most to prediction. B) Precision-Recall curve for prediction of severity classes. C) ROC curve for prediction of symptom severity. D) SHAP score of each feature.

**Table 5:** Accuracies of multiple models for Predicting severity of individual using Binary Prediction.

| Sr. No. | Classifier | Accuracy | F1-Score |
| --- | --- | --- | --- |
| 1. | Xgboost | 0.9594 | 0.9401 |
| 2. | Neural Network | 0.9864 | 0.9753 |
| 3. | Decision Tree | 0.9459 | 0.9631 |
| 4. | Logistic Regression | 0.9661 | 0.9563 |
| 5. | Random Forest | 0.9729 | 0.9842 |
| 6. | Ensemble Learning | 0.9913 | 0.9941 |

**Table 6:** Accuracies of multiple models for predicting classes of severity using Multiclass prediction.

| Sr. No. | Classifier | Accuracy | F1-Score |
| --- | --- | --- | --- |
| 1. | Xgboost | 0.7586 | 0.7364 |
| 2. | Neural Network | 0.7613 | 0.7745 |
| 3. | Decision Tree | 0.5517 | 0.5067 |
| 4. | Logistic Regression | 0.5287 | 0.5265 |
| 5. | Random Forest | 0.5919 | 0.6037 |
| 6. | Ensemble Learning | 0.7873 | 0.7902 |

## 5 DISCUSSION

The main objective behind this work was to determine a common source of variability between ASD and typically developing to identify heterogeneity within ASD samples that shows different ToM functional connectivity. We identified ToM functional connectivity brain features that distinguish individuals with ASD from TD control subjects and accurately predict social cognition impairment, predicting clinical symptom severity using a novel end-to-end Ex-AI model. Further to this we hypothesized that after controlling for a common source of variability in ASD and TD samples, the distinguishable SFC ToM features could be used to train the model to learn the connectivity patterns among distributed brain areas primarily anchored in ToM and Default Mode brain regions without ad hoc feature engineering, achieving high classification accuracies in cross-validation analysis of data from the multisite ABIDE cohort. This approach overcomes the classification limitation using SFC previously employed by many researchers. Crucially, our results demonstrate that resting-state FC from Theory-of-Mind and DMN brain regions can distinguish between ASD and typically developing and address several existing challenges. Feature identification using an IG, LIME, and Shap approach revealed that brain features associated with the key nodes of the ToM, DMN, and cognitive control systems (ref) most clearly distinguished ASD from TD control subjects in the ABIDE cohorts and the ToM nodes predicted core social and communication deficits, but not restricted and repetitive behavior (RRB), phenotypic features associated with ASD.

This study’s first challenge was identifying Theory-of-Mind brain regions so that functional connectivity between them could be used as the best feature set for classification. We extracted a total of 16 (12 ToM regions and 4 other regions associated with ToM) brain regions associated with the Theory-of-Mind and Default-mode network and extracted Bold time-series signals [5]. After calculating Pearson’s correlation analysis, we generated a static functional connectivity(SFC) matrix and represented it in a Low-dimensional embedding. We found hyper-connectivity within the TOM network in the ASD group (Left TPJ, Right TPJ, Posterior cingulate cortex (PCC), Precuneus, Left angular gyrus, Right angular gyrus, and Right superior temporal sulcus), whereas hypo-connectivity within the TOM region in the TD control group in the resting state. In the previous study, the author also reported hyperactivity in autism compared to typically developing in activity and resting state [35]. This study’s second challenge was finding a common source of variability that may find overlapping connectivity patterns across distributed brain regions. To this end, we used the CVAE model containing encoding and decoding processes to generate latent space and RSA methods to calculate the dissimilarity matrix; Subsequently, the Kendall correlation coefficient showed that ASD-specific characteristics were consistently correlated with clinical measures like ADOS-Total scores, ADOS-Social scores, FIQ, and resting-state functional connectivity. In contrast, shared variation was associated with non-clinical measures like Age, Gender, and Scanner Id. Results of the CVAE model demonstrated that segregating ASD-specific variations from shared variations revealed individual differences and discovered new feature sets for training the classification model. The previous study [26] reported that the CVAE model could not find any sub-groups within ASD samples using structural brain images. The current study extended the previous approach by employing a similar methodology on resting-state functional scans based on selecting specific ToM and DMN brain regions. Our results based on the CVAE model could not identify specific hidden subgroups within ASD samples; however, the model could robustly identify non-overlapping features between ASD and TD.

Next, we developed a novel, explainable and interpretable ML/DNN (ExML/DNN) model to distinguish between ASD and typically developing. The proposed ExDNN and ExMl models identified the ASD samples and achieved, on average, 95 % accuracy with an F1 score of 0.94. These models surpass the conventional approaches that use SFC and Bold time-series data from complete brain parcels [28], [36], [37], [38]. Our approach suggests that resting-state functional connectivity from Theory-of-mind brain and associated DMN brain regions could be useful feature sets that can distinguish ASD from typically developing accurately. To validate the results, we performed five-fold cross-validation and a leave-one-out approach. From five-fold cross-validation, We were able to get, on average, 97 % accuracy with an F1-score of 0.96, and from leave-one-out, we got 96 % accuracy.

The fourth limitation of the study was to Identify neuro-biological features that were associated with ASD samples and that underlie classification. Conventionally, deep learning and machine learning models are trained by contriving machine-generated intermediary feature sets, which can not be explainable by humans [39], [40], [41]. In such a case, there is a need to know human-interpretable features that underlie classification and their associative importance [42], [43], [44]. Here, we used Integrated Gradient (IG) for deep learning; we applied SHAP and LIME for machine learning models that developed an automated AI method for identifying neuro-biological features and hyper-parameters and ranked them in order of importance [45], [46], [47], [48]. The framework we have proposed here suggests that functional connectivity between Left temporoparietal junction, Right temporoparietal junction, Right superior temporal sulcus, Left angular gyrus, Posterior cingulate cortex, and precuneus brain regions are the most robust feature set and could be potential biomarkers during early development that distinguish ASD from typically developing.

Given the heterogeneity and extent associated with ToM in ASD, the fifth challenge we addressed was to uncover neurobiologically interpretable ToM connectivity subtypes and features associated with the severity of social and communication deficits, a core defining characteristic of the disorder. This study addressed the above challenge by identifying sub-groups within the ASD samples using resting-state SFC. We used the K-means clustering algorithm, identifying three hidden sub-groups that showed inter-cluster differences. Moreover, we identified the same three clusters using a variational autoencoder model with K-means. Thereafter, we estimated SFC between ToM and DMN regions for all sub-groups and found that the functional connectivity in the first sub-group resembled typically developing controls, i.e., displaying hypo connectivity. In contrast, we found hyper-connectivity in the other two subgroups. Subsequently, We calculated the difference in functional connectivity patterns for each group and discovered that all sub-groups were distinct.

This study’s final and sixth challenge was identifying neurobiological features associated with ASD social and communication deficits and predicting ADOS-Total and ADOS-Social symptom severity scores. The results obtained from the previous challenge indicated that the first sub-group was closely associated with TD. To contextualize the above result, using resting-state functional connectivity, here we defined four classes of severity: low, moderate, high, and very high, and applied Explainable machine learning and deep learning algorithms to predict the severity of each individual. We obtained 76 % accuracy using Ex-DNN. Next, we applied Connectome-based prediction modeling (CPM) to predict the ADOS-Total and ADOS-Social scores of each individual from the ASD samples [34], [49]. We extracted time-series signals from Theory-of-Mind and Default-mode networks, calculated each individual’s functional connectivity (FC) matrix, and fitted behavioral scores and FC to a linear model to predict scores. The CPM model could predict ADOS-Social scores accurately but not ADOS-Total crucially, suggesting the specificity of the ToM brain connectivity features. This further suggests that whole brain SFC among distributed brain regions encompassing sensory, subcortical, and heteromodal higher-order cortical brain regions may be necessary to predict ADOS-Total symptom severity scores accurately. Overall, the brain inspired framework proposed here accurately distinguished ASD from control subjects and uncovered distinguishing neurobiological features of ToM and DMN functional brain patterns. Our discovery also provides unique predictive fingerprint in each individual subjects in ASD group that robustly predicted their severity of social and communication deficit.

## 6 CONCLUSION

Our End-to-End AI approach was able to distinguish the ASD samples from typically developing on multisite ABIDE cohorts using Theory-of-Mind brain regions with high accuracy. The idea behind disentangling ASD-specific variation from shared variation revealed correlations between individual differences without the need for any additional training. Our proposed Ex-AI approach using Theory-of-Mind brain regions for understanding Social cognition deficits in ASD subtypes overcomes key methodological challenges in classification. It addresses an even deeper problem concerning explainable and interpretable biological features offered by existing AI models of ASD social symptom severity. Our approach not only classifies ASD samples but also identifies brain features associated with ASD that underlie the classification. It also identifies heterogeneity or sub-groups within ASD samples and accurately predicts each individual’s mild to severe social symptom severity and open up future possibilities of individual fingerprinting. There are two major limitations of our study. First, we did not carry out any out of the sample generalization as we focus exclusively on ABIDE cohort for developing our framework and secondly, our methods were not tested with cognitive task associated with ToM data from ASD and TD individuals. However, some work along this line is already underway in our lab and will be reported elsewhere. Taking together our discovery of robust individualized functional brain connectivity biomarkers of relevant ToM regions in ASD could transform our understanding of the etiology, diagnosis, and trait specific modulation of connectivity in pervasive neurodevelopmental disorder. More generally, our approach provides new Ex-AI-based tools for probing the robust and interpretable neurobiological bases of other neurodevelopmental and psychiatric disorders and the underlying clinical symptoms, with the potential to inform precision neuroimaging in brain disorders.

## 7 ACKNOWLEDGMENT

We acknowledge the generous support of IIT Jodhpur Core funds and the Computing facility. D.R. acknowledges the generous support of the NBRC Flagship program BT/MEDIII/NBRC/Flagship/Pro-gram/2019: Comparative mapping of common mental disorders (CMD) over the lifespan.

## Supplementary Methods

### ABIDE Dataset

In the current study, we used neuroimaging and phenotypic data from ABIDE I dataset comprised of 400 Autism and 460 neuro-typicals samples from 20 sites (Total no. of samples per site: Mean = 43, range: 39-71).

### fMRI Preprocessing

The neuroimaging functional images were preprocessed using Configurable Pipeline for the Analysis of Connectomes (C-PAC) based on AFNI, ANTs, FSL, and python code. Subjects with poor-quality data were excluded from the final analysis (Total no. of images = 952, After preprocessing: total no. of images = 860). Prior to data contribution to ABIDE I all the sites (including data used in this study) were required to confirm clearance from their local institute review board (IRB) or ethics committee. The local ethics committee has approved both the initial data collection and the retrospective sharing of a fully de-identified version of the datasets (i.e., after removal of the 18 protected health information identifiers including facial information from structural images as identified by the Health Insurance Portable and Accountability Act [HIPAA]). All study participants provided written informed assent or consent prior to study participation with parental or legal guardian consent required of all study participants under the age of 18. All the fMRI images were reorganized for head motion correction to the average time frame using AFNI’s 3dvolreg, and slice time correction was performed using AFNI’s 3dTshift. The global mean intensity was normalized to 10000. We applied nuisance signal regression that included motion parameters and linear and quadratic trends and also applied band-pass filtering from 0.01-0.1Hz, which helped to remove low-frequency artifacts. But band-pass filtering was not able to remove time-varying means and covariances artifacts. To address this issue, registered the functional images in anatomical space using a linear transformation (see Supplementary material section 1 for details).

As we were dealing with data from multiple sites, there was the issue of heterogeneity. To address this issue, we applied the combat harmonization technique, that harmonized age, gender, and scanner id-related heterogeneity. It took all these parameters as input in .csv file format and performed harmonization using multivariate linear mixed-effects regression. This method used empirical Bayes for the improvement of estimation of the model.

### Resting-State Functional Connectivity

Theory-of-mind (ToM) is the concept of understanding others’ beliefs, desires, intentions, and emotions that influences social interaction. Previous studies have shown that the frontal posterior network has been activated during the Theory-of-Mind task that included medial prefrontal cortex (MPFC), Posterior cingulate cortex (PCC), and Bilateral temporoparietal junction (LTPJ and RTPJ). These core brain areas deal with theory-of-mind, but some other brain regions like right anterior superior temporal sulcus medial precuneus are also activated during TOM-related processing. On the other side, individuals with Autism mainly face the problem of social and communication deficits and repetitive behavior, i.e., a deficit in ToM.

In the current study, we extracted activation time series BOLD signals from Theory-of-Mind and other brain networks associated with social cognition, i.e., Default Mode Network(DMN) as listed in Table 2 below. The Regions-of-Interest(ROI) for this study was derived from the Human Connectome Project(HCP) available on Neuro Vault. We used the Harvard-oxford atlas and created ROIs using a spherical binary mask with a 10 mm radius. To validate the results, we performed multiple one-sample t-tests with a p-value<0.01 and applied FDR correction.

To check within network and between network connectivity from growing age, we also stratified age into 3 groups: 6-12 yrs, 13-18 yrs, and 19-25 yrs. We calculated the functional connectivity matrix for all the groups using ToM and DMN areas. We found that in 6-12 yrs group ToM areas (i.e., LTPJ, RTPJ, MPFC, PCC) were highly connected in ASD group, but this condition was not true in TD group, whereas we were not getting hyper-connectivity in DMN areas for the same group. For the second group i.e., 13-18 yrs, we found hyper-connectivity within ToM network in ASD group, whereas hypo-connectivity in TD group. When we included third group 19-25 yrs, we observed hyper-connectivity in TD group and hypo-connectivity in ASD group.

**Figure 1:**
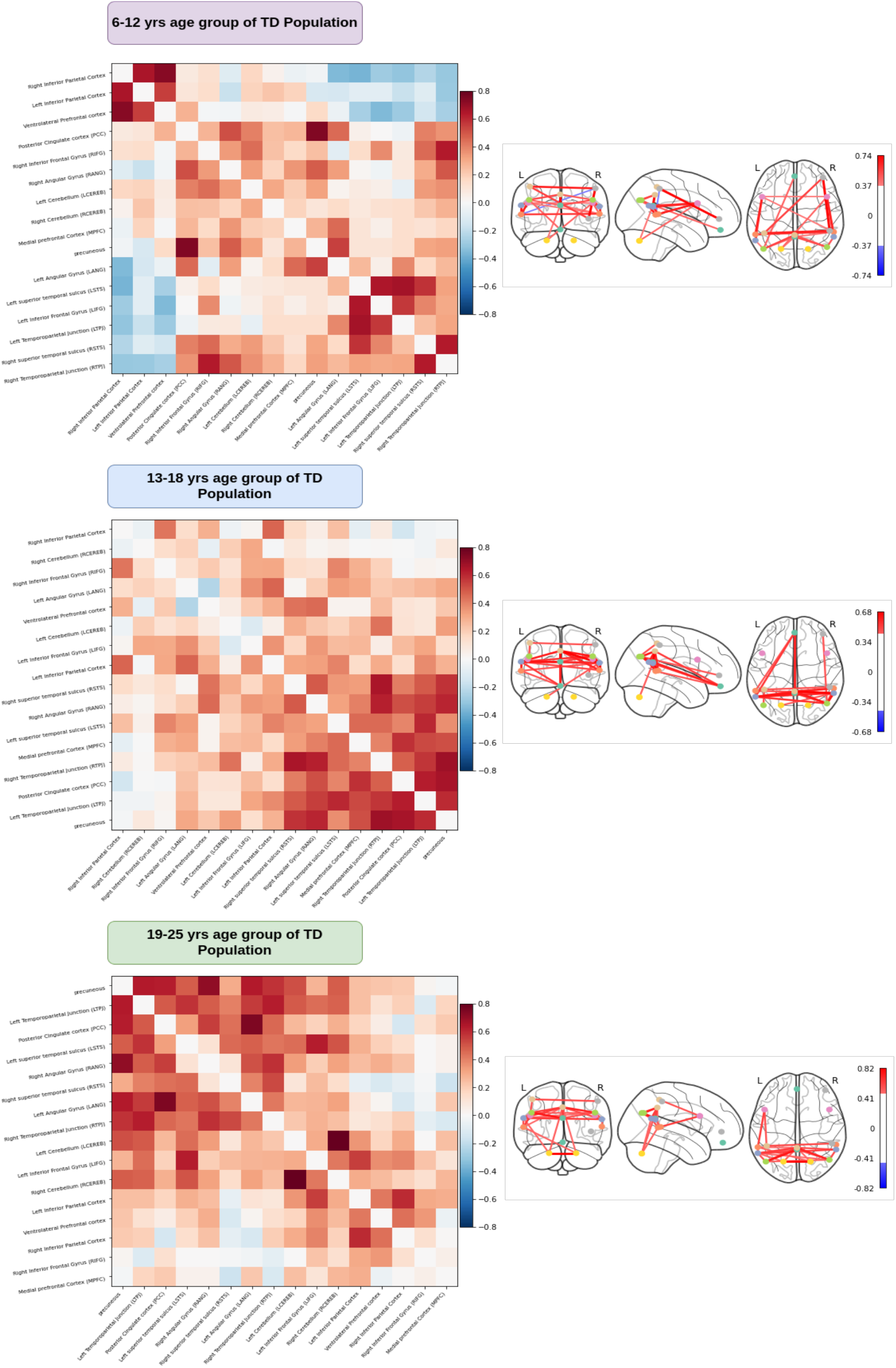
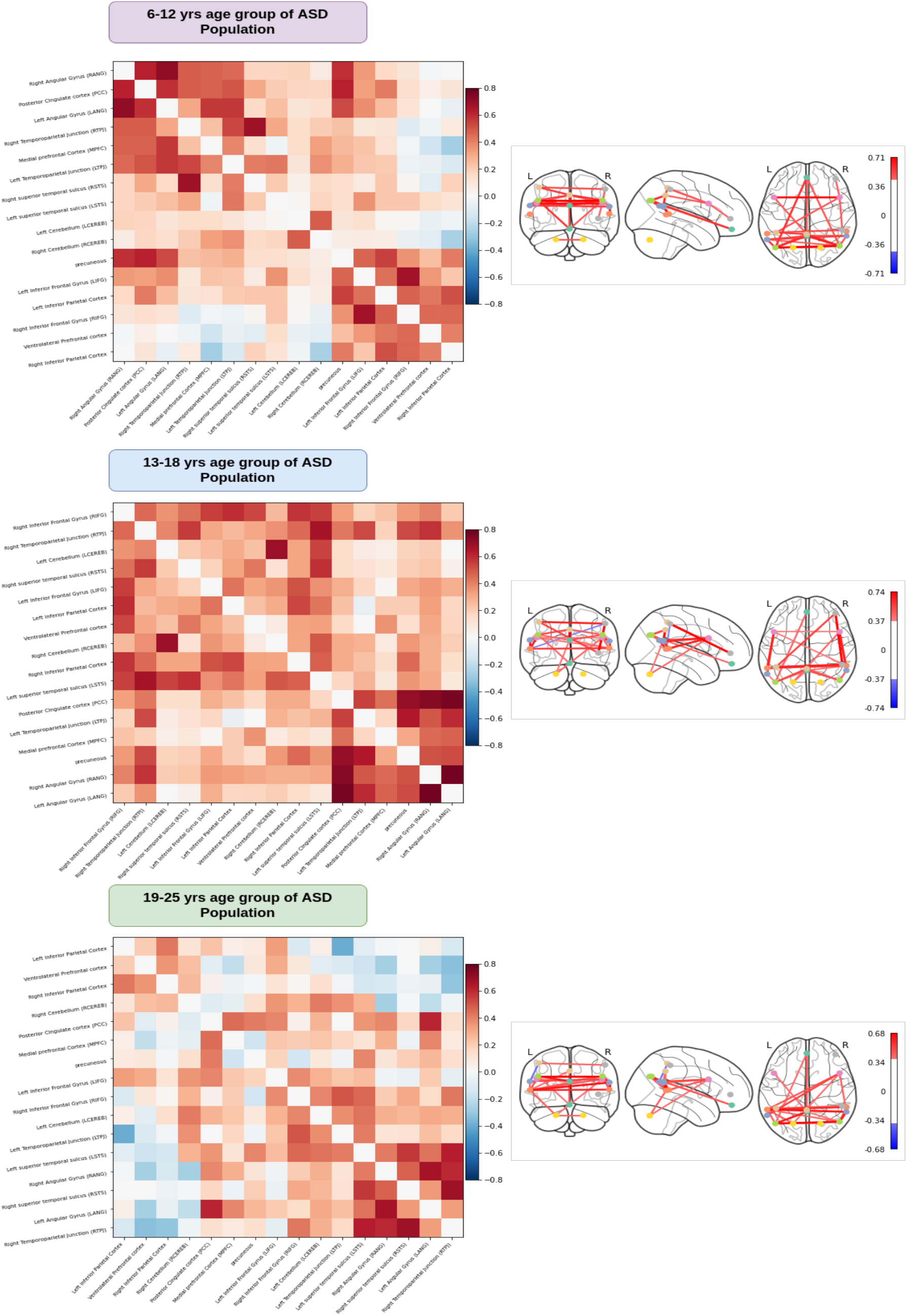
Functional connectivity matrices for different age group for ASD and neuro-typical population.

### Identification of Shared and Specific Variability

We hypothesized that common source of variability between neuro-typicals and ASD individuals could be identified using what was common b/w TD & ASD groups, Whereas what was not common in both the groups that differences could be the best possible characteristics to identify features on which the classification model could be trained to learn characteristics difference and be able to classify ASD & TD given some test dataset & also able to predict their association with the symptom severity scores on behavior. To identify ASD-Specific variation, we disassociated ASD-specific variation from common variation between Neuro-typical & Autistic using a Contrastive-Variational Autoencoder model (CVAE) that allowed us to distinguish clinically applicable individual variationWe found that shared variation was associated with non-clinical characteristics (i.e, Scanner Id, Age, Gender), whereas ASD-specific variation was related to clinical measures(ADOS-Total, ADOS-Social, DSM-IV, FIQ) as well as resting-state functional connectivity.

### Classification Analysis Using Five-Fold Cross Validation

To reduce the chance of bias and report low variance, we implemented five-fold cross-validation to evaluate model’s performance (precision, recall, accuracy, F1-score). We divided the complete dataset into five parts, which used four parts for training and validation purposes and five-part for testing. We repeated this process five times for each iteration. We then calculated the average accuracy and F1 score. We also implemented a leave-one-out approach, in which we divided the dataset into two parts training and testing, trained the model on the complete training set except for the leave-out observation, and performed testing on the leave-one-out observation. We repeated this process n-times for each iteration and reported the average accuracy.

### Identification of Sub-Group Within ASD Group and their Functional Connectivity Differences

After performing classification, we were able to identify the ASD samples. Resting-state functional connectivity from ToM and DMN areas were provided to an unsupervised k-means algorithm based on Euclidean distance to identify heterogeneity or subgroups within ASD population. Using the elbow approach, we obtained three sub-groups of different sizes. We also performed validation of results using variational autoencoder model, in which applied K-means on latent space and again got 3 clusters.

**Figure 2:**
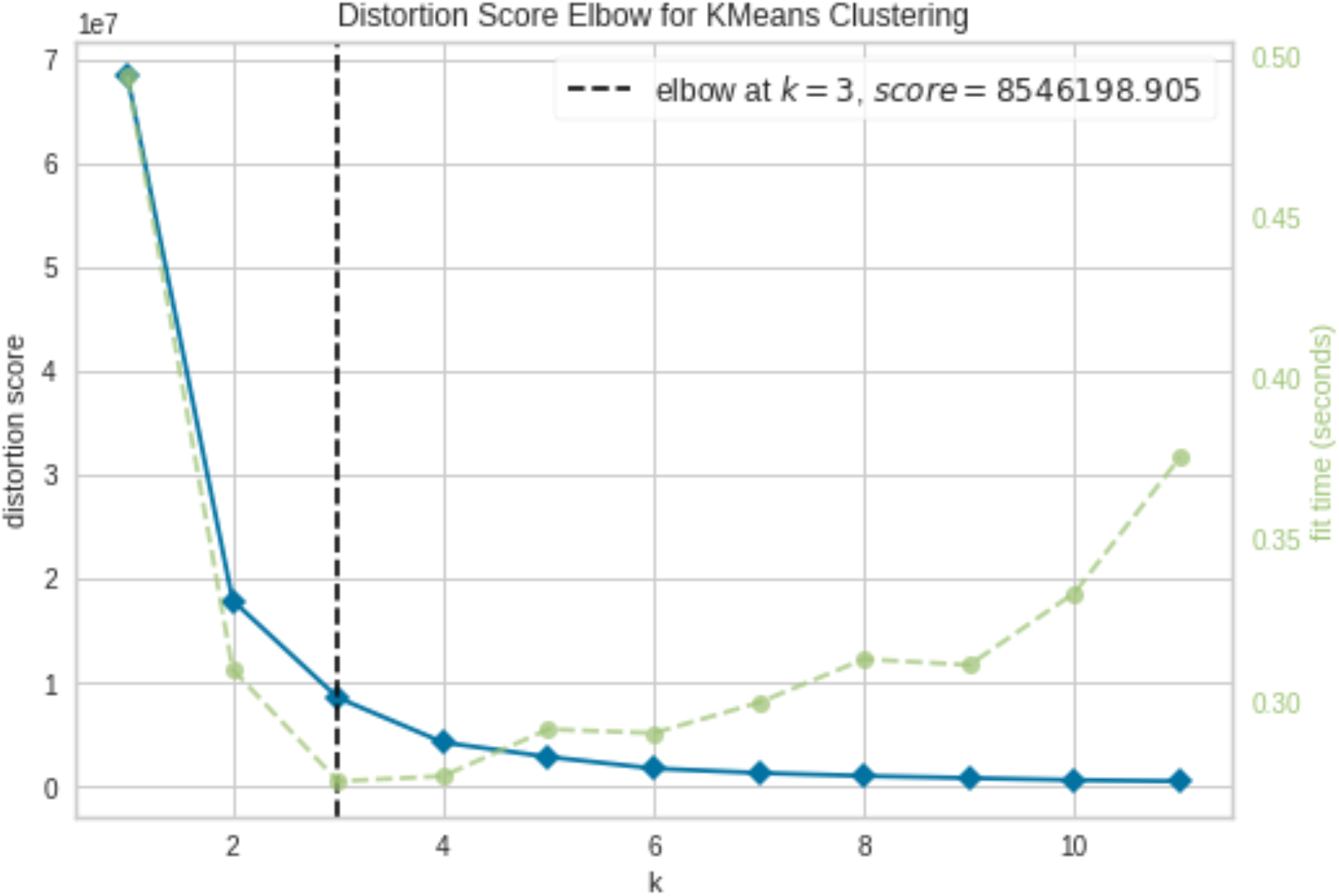
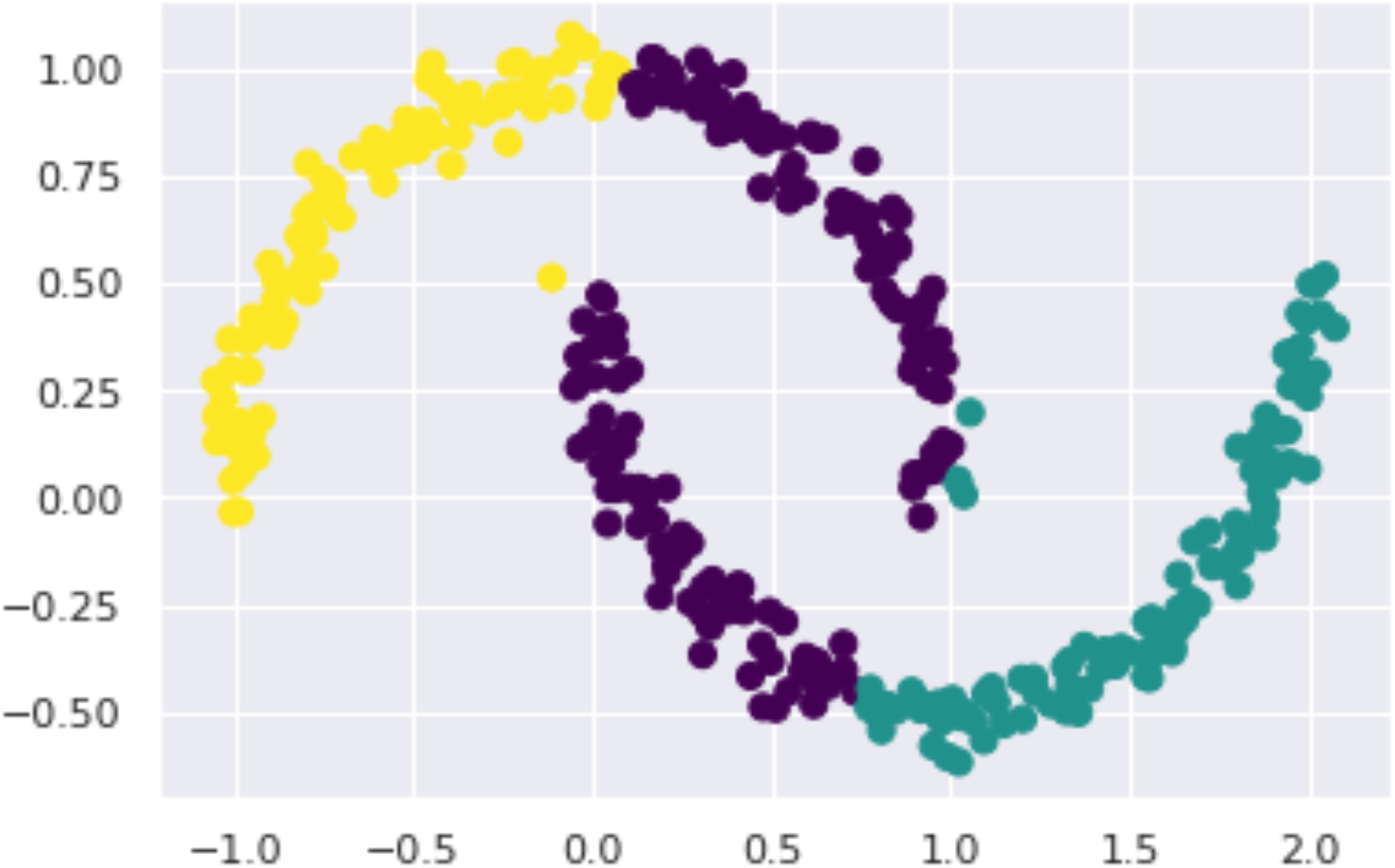
K-means clustering approach to identify sub-groups within ASD Population. A) Elbow approach to identify no. of clusters, b) Visualization of each cluster.

## Supplementary Results

### Identification of Shared And ASD-Specific Features or Source of Variability

To recognize the source of variability between ASD and neuro-typicals, we first implemented the Variational Autoencoder model(VAE) as a base model with a single latent space that showed correlation with some non-clinicalcharacteristics (ScannerID (τ = 0.01, t(9)=0.74, p-value < 0.042), Gender (τ = 0.01, t(9)=1.94, p-value < 0.04)) and also correlated with DSM-IV(Diagnostic Statistical Manual IV)(τ = 0.02, t(9)=0.54, p-value < 0.03) but did not get any correlation with ADOS Total (τ = -0.01, t(9)= -2.96, p-value < 0.031), ADOS Social (τ = -0.01, t(9)=-2.66, p-value < 0.026), FIQ (τ = 0.00, t(9)= -1.37, p-value < 0.027), and functional Connectivity (τ = 0.00, t(9)=-5.56, p-value < 0.04). Hence, the VAE model failed to encapsulate variation in ASD and TD. To overcome the limitation of VAE, we implemented the Contrastive Variational Autoencoder model(CVAE). As a result, we found that shared characteristics were correlated with non-clinical characteristics (i.e., ScannerId (τ = 0.01, t(9)=0.85, p-value < 0.0.39), Age (τ = 0.03, t(9)=1.41, p-value < 0.04), Gender (τ = 0.02, t(9)=0.51, p-value < 0.045)); in contrast, ASD-specific characteristics were associated with clinical measures (i.e., ADOS Total (τ = 0.04, t(9)=2.65, p-value < 0.03), ADOS Social (τ = 0.04, t(9)=1.88, p-value < 0.01), FIQ (τ = 0.02, t(9)=1.95, p-value < 0.04)), as well as Functional Connectivity (τ = 0.04, t(9)=1.32, p-value < 0.02).

### Classification of ASD and Neuro-typical Individuals Using Explainable AI

Our ExAI model outperformed the traditional approaches that use a) Functional connectivity matrix as a feature set, which achieved an average accuracy of 55% with an F1 score of 0.52. In this work, we randomly selected samples with a ratio of 70:30 for training and testing sets. Then we trained our models in various scenarios: a) We trained our model on shared characteristics set that achieved an average accuracy of 60% from ExDNN with an F1 score of 0.58 and 80% from ExML with average F1 score of 0.81. b) then, the models were trained using shared and ASD-specific characteristics that achieved an average accuracy of 85% using ExDNN with an average F1 score of 0.79 and average accuracy of 90% with an F1 score of 0.88 using ExML. c) Finally, we trained our models with resting state FC of Tom and DMN areas as feature set and achieved average accuracy of 96% with F1 score of 0.95, sensitivity of 0.95, and specificity of 0.93 using ExDNN model, and average accuracy of 90% with average F1 score of 0.91 and sensitivity of 0.87 and specificity of 0.85 using ExML. To validate the results, we also performed five-fold cross-validation and a leave-one-out approach and achieved average accuracy of 96% using ExDNN and average accuracy of 97% using ExML.

We performed classification on different age groups (i.e., 6-12, 13-18, and 19-25 yrs age group): we trained our model on different age groups and test it on different age groups. We repeated this process for all age groups. We were getting average accuracy of 93 % using ExDNN with F1 score of 0.91 and average accuracy of 89 % using ExML with F1 score of 0.87. We observed that connectivity between Left temporoparietal junction, Right temporoparietal junction, Right superior temporal sulcus, Left angular gyrus, Posterior cingulate cortex, and precuneus brain regions contributed the most in classifying ASD samples.

**Figure 3:**
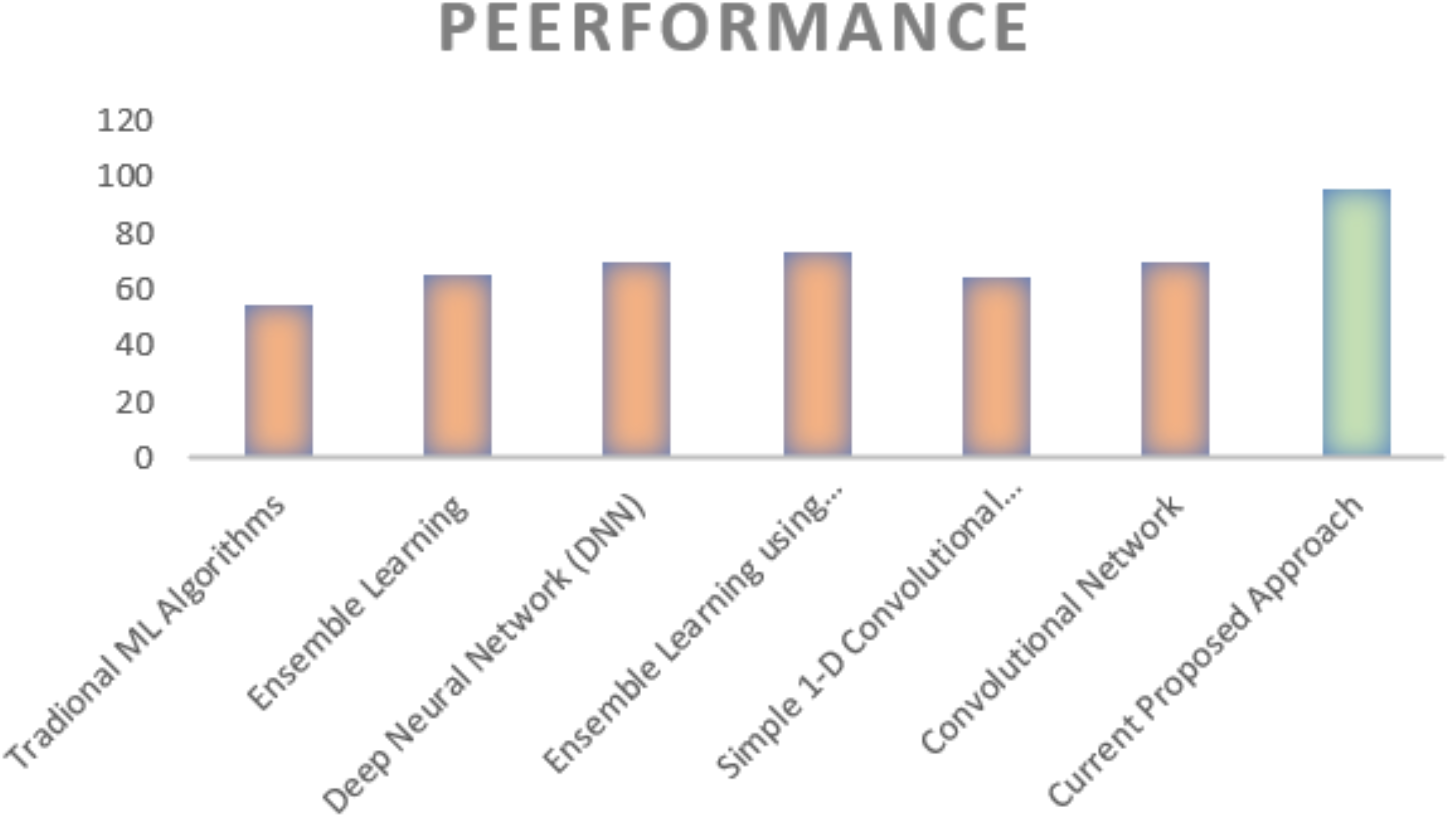
Previous studies and their performance using whole brain on ABIDE cohort.

**Figure 4:**
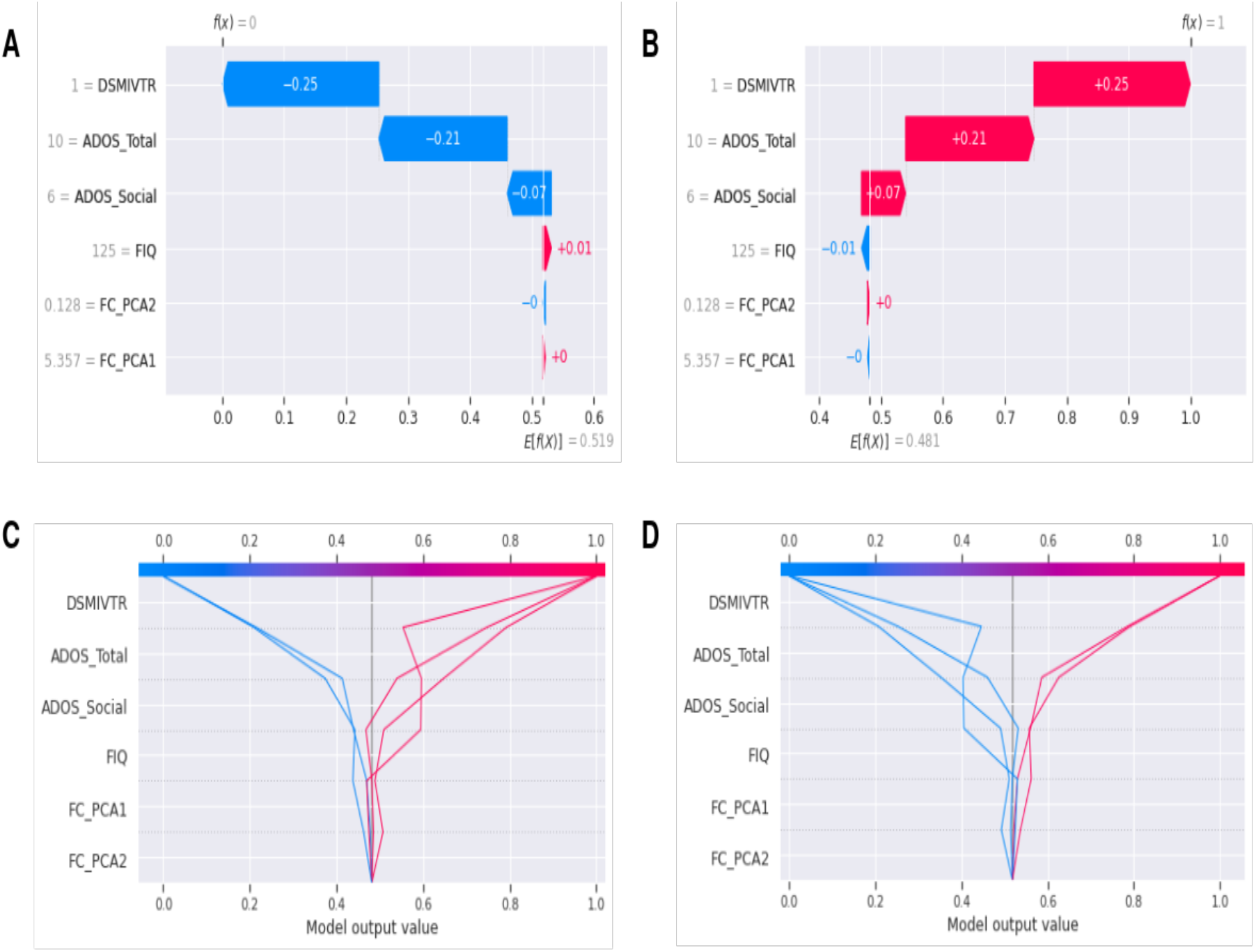
SHAP values for each features that underlie classification between ASD and Neuro-topicals.

### Symptom-Severity Score Prediction

To check the severity of each individual, we implemented binary prediction using multiple Explainable AI algorithms (ExAI) on ASD-population and achieved an average accuracy of 96.5% with F1 score of 0.97. After that, to predict classes of severity, we implemented multi-class prediction in which we divided classes into four parts: Low, Moderate, High, and Very High, and achieved an average accuracy of 67% with F1 score of 0.66. We implemented Connectome-Based Prediction modeling (CPM) to predict individual behavior scores. We were getting better results in ADOS-Social score prediction. To validate the results, we performed five-fold cross-validation that achieved an average accuracy of 90% with F1 score of 0.92.

## Notes

### Competing Interest Statement

The authors have declared no competing interest.

